# Membrane Shape Remodeling by Protein Crowding

**DOI:** 10.1101/2020.10.22.351130

**Authors:** Susanne Liese, Andreas Carlson

## Abstract

The steric repulsion between proteins on biological membranes is one of the most generic mechanisms that cause membrane shape changes. We present a minimal model where a spontaneous curvature is induced by steric repulsion between membrane-associated proteins. Our results show that the interplay between the induced spontaneous curvature and the membrane tension determine the energy minimizing shapes, which describe the wide range of experimentally observed membrane shapes, *i*.*e*. flat membranes, spherical vesicles, elongated tubular protrusions, and pearling structures.

## Introduction

Membrane nanotubes formed by lipid bilayers are ubiquitous in cell biology [1, 2] where they play a key role in various transport processes by facilitating the inter- and intracellular transport of fluids and macromolecules [2, 3]. The high surface-to-volume ratio inherent to tubular shapes enables rapid exchange of biomaterial across the lipid bilayer by transmembrane proteins or protein channels [4, 5]. As membrane tubes can bridge distances of several micrometers, they can facilitate the transport of nutrients or even entire organelles such as mitochondria and lysosomes between cells [6, 7].

A wide range of biophysical mechanisms cause tube formation, such as the adsorption of intrinsically curved proteins [8, 9, 10, 11], internal and external protein scaffolds [12, 13], local pulling forces [14], membrane compression [15], osmotic deflation [16, 17], and protein crowding [18, 18, 19]. Protein crowding, *i*.*e*., the accumulation of proteins in a confined membrane domain is a phenomenon that is ubiquitous in biological membranes, which typically contain a multitude of domains with densely packed proteins of different size and where an over-expression of specific proteins or bio-polymers can be a disease marker. Tumor cells, for example, exhibit both a higher concentration of glycosylated polymers and a higher tendency to form tubular protrusions [20], indicating that crowding induced membrane remodeling sets healthy and malignent cells apart.

Despite that protein crowding plays an essential role in many membrane remodeling phenomena, a theoretical model encompassing the protein size and density that accurately predicts the wide range of membrane shapes observed experimentally is still lacking. Theoretical studies have shown that the steric pressure between dense, unstructured proteins as well as an asymmetry in the protein distribution across the lipid bilayer produces a spontaneous membrane curvature [21, 22, 23] and thus induces changes in the membrane shape. Stachowiak et al. observed membrane tubes protruding from synthetic giant unilaminar vesicles (GUVs) functionalized with green fluorescent proteins (GFPs) [24]. A schematic description of this experimentally observed tubulation process, which we theoretically investigate here, is shown in Fig. 1a [24]. The membrane tubes completely consume the GFP-coated membrane region (shown in blue in Fig. 1a) and extend over a length of several micrometers, which is of the same order of magnitude as the GUV diameter [24]. Other polymers containing a large unstructured region also generate a repulsive effect that leads to tube formation [20, 25, 26]. Moreover, Shurer et al. recently demonstrated that densely packed brush-like glycocalyx polymers induce a variety of cell membrane shapes; flat membranes, Ω-shaped membranes (referred to as blebs), tubes and pearls, induced by densely packed brush-like glycocalyx polymers, as shown in Fig. 1b [20]. The wide range of polymers causing membrane tubes, including folded, intrinsically disordered and brush-like polymers [20, 24, 25], suggests that the shape remodeling process does not depend on the chemical structure of the individual protein, but is based on a generic mechanism that is potentially relevant for a variety of proteins in densely packed membrane domains. However, a biophysical model that reveals the minimal requirements for the formation of the different and seemingly incommensurable membrane shapes through a single mechanism has yet to be established. To study the biophysical origin of the different membrane shapes observed experimentally, we derive a minimal mathematical model and describe the progressive shape transformation from a flat membrane to a fully formed tube induced by protein crowding.

**Figure 1:**
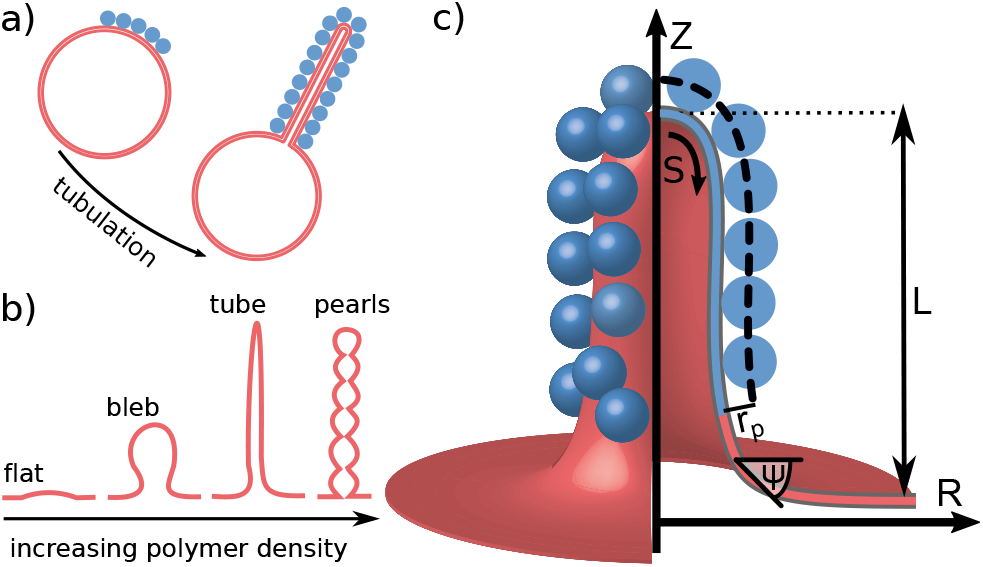
a) Schematic illustration of the membrane tube formation as observed experimentally on GUVs [24]. The protein coated domain (shown in blue) is completely consumed by the protruding tube. The size of the vesicle (red) and the tube (blue) are drawn to scale to illustrate a ratio between the protrusion height *L* and the crowded domain *A*_c_ that is representative of the order of magnitude found in experiments [24]. b) Qualitative representation of membrane shapes observed experimentally by Shurer et al. [20] on cells with glycocalyx biopolymers with increasing polymer density. c) The membrane shape is parameterized by the arc length *S* and the azimuthal angle *ψ*, where we treat the membrane as axially symmetrical around the *Z* axis. The proteins that bind to the membrane within an area *A*_c_ (shown in blue) are modeled as spheres with a radius *r*_p_.

## Mathematical Model

We start by formulating the membrane energy *E*, which takes into account the Helfrich bending energy [27, 28], the membrane tension *σ* and the lateral pressure *p* between crowded proteins:

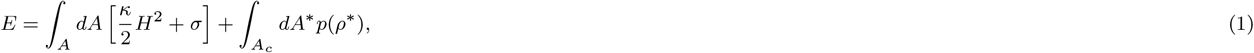

with *A* the total membrane area, the bending rigidity *κ* and the mean curvature *H*. We refer to the area *A*_c_, in which proteins are bound to the membrane, as the crowded domain. The proteins are modeled as spheres with radius *r*_p_ and density *ρ*, which experience the dominant contribution of steric repulsion along a virtual surface shifted by *r*_p_ perpendicular to the membrane (dashed line in Fig. 1c). The variables related to the shifted surface are indicated by an asterisk (*) in Eq. 1. In the model presented here the proteins adsorb to one side of the lipid bilayer. A similar energy expression is obtained for transmembrane proteins, where *r*_p_ would correspond to the size of the protein portion protruding from the membrane. We consider a crowded domain much smaller than the total area, which means that also the volume of the protruding shape is small compared to the total volume of the vesicle. Therefore the associated transmembrane pressure has no influence on the membrane shape, see the Supplemental Material (SM). Experimental studies have shown the simultaneous formation of multiple protrusions [20, 26], which might arise due to heterogeneity in the protein distribution or within spatially separated crowding domains. Since the tubulation mechanism is the same within each crowding domain, we limit the model presented here to the formation of a single protrusion.

The membrane shape is parameterized by the arc length *S* and the azimuthal angle (Fig. 1c), where we model the membrane as a thin axially symmetric surface, which is valid as long as the radius of curvature is large compared to the membrane thickness [28]. The height *Z* and the radial coordinate *R* are then obtained via *dR/dS* = cos*ψ* and *dZ/dS* = − sin *ψ*. If the protein radius *r*_p_ is small compared to the two principle curvatures, *C*_1_ = *dψ/dS* and *C*_2_ = sin *ψ/R*, the area element and the density along the shifted protein surface can be expressed as *dA*^∗^ = *dA*(1 + *r*_p_*C*_1_)(1 + *r*_p_*C*_2_) and *ρ*^∗^ = *ρ/*[(1 + *r*_p_*C*_1_)(1 + *r*_p_*C*_2_)] (see SM). The lateral pressure *p* = *p*(*ρ*^∗^) depends on the membrane curvature through the curvature dependence of *ρ*^∗^, where the relationship between the lateral pressure and the protein density is expressed in the most general form as a power series by a virial expansion 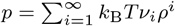, with the thermal energy *k*_B_*T* and the virial coefficients *v*_*i*_. Inserting these relations into Eq. 1 and expanding up to second order in *C*_1_ and *C*_2_, we write the membrane energy up to a constant as the sum of two terms, *i*.*e*. the bending energy *E*_*κ*_ and the tension energy *E*_*σ*_ :

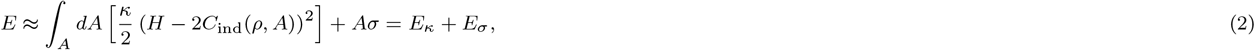

where the influence of the lateral pressure between the proteins is expressed by an induced spontaneous curvature *C*_ind_. We consider a homogeneous protein distribution (*ρ* = *const*.) within the crowded domain *A*_c_ and *ρ* = 0 in the protein free domain. In general, proteins can diffuse within the membrane, which then can cause a spatial variation in the spontaneous curvature. However, heterogeneity of the protein or lipid distribution is counteracted by an energetic penalty ∼ (∇*ρ*)^2^ that maintains a more uniform density [19, 29]. To keep the model minimal, we consider only the limit of small density gradients, assuming *ρ* = *const*. and *C*_ind_ = *const*..

We find the following expression for the induced spontaneous curvature:

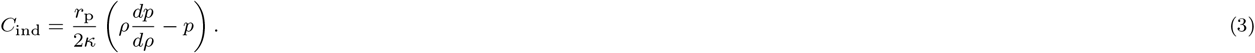

A detailed derivation of Eqs. 2, 3 is presented in the SM. To derive the relation between the induced spontaneous curvature and the lateral pressure (Eq. 3), we take into account a spatially varying membrane curvature. Eq. 3 thus provides a more general version of the result previously obtained by Derganc *et al*., who find a linear relationship between the induced curvature and the lateral pressure *p* under the constraint of a uniform membrane curvature [21]. If the steric repulsion of proteins is caused by volume exclusion alone, the lateral pressure is well approximated by the Carnaham-Starling equation [30]:

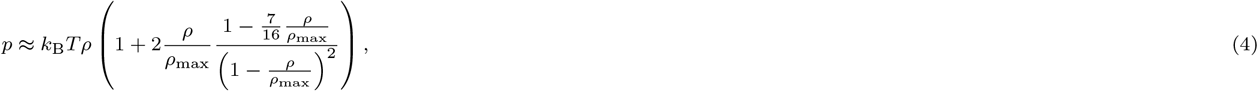

where the maximal areal density is set by the protein radius with 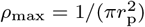. Hence, the induced spontaneous curvature reads:

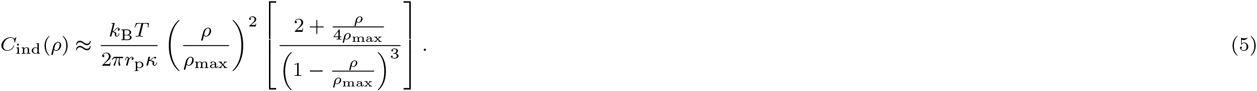

The prefactor in Eq. 5 shows an inverse scaling of the induced spontaneous curvature with respect to the size of the proteins *r*_p_ in accordance with previous theoretical studies [21, 22]. In addition, Eq. 5 provides an explicit relation between *C*_ind_ and the scaled protein density *ρ/ρ*_max_.

We find that the lateral pressure term in Eq. 1 also causes tho additional energy terms that only act in the crowded domain; an increase in bending stiffness Δ*κ* and a decrease in Gaussian bending rigidity *κ*_g_ (see SM, Eq. S23). The additional contributions to the bending rigidities Δ*κ* and *κ*_g_ scale with 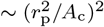 and are therefore negligible, since the crowded domain is large compared to the size of the proteins.

We use the following parameters typical to experimental systems; *κ* = 200k_B_T [31], *σ* = 10^−6^N/m [32, 33], *A*_c_ = 25*µ*m^2^ [24], unless otherwise specified. The scaled membrane tension then becomes *σA*_c_*/κ* = 25.

When minimizing the energy (Eq. 2) the membrane can take two characteristic shapes; a rather flat shape that minimizes *E*_*σ*_ or a cylindrical shape that follows the induced spontaneous curvature and minimizes *E*_*κ*_. Before we derive the shape equations, we discuss two analytical approximations in the limit of small and large *C*_ind_ to gain an intuitive understanding of the scaling of the protrusion height *L* and the energy. For small *C*_ind_ we describe the crowded domain as a spherical cap with a radius *R*_s_ and an opening angle *α*, while the protein-free membrane region is considered to be flat, see Fig. 2a. The energy difference Δ*E* between a flat and a deformed membrane according to Eq. 2 is 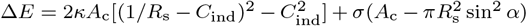. The radius *R*_s_ and the angle *α* are related via the area as 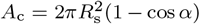. Minimizing Δ*E* with respect to *R*_s_ for fixed *A*_c_ and *C*_ind_ results in *L* = *A*_c_*C*_ind_*/*[2*π* + *σA*_c_*/*(4*κ*)] (dashed line in Fig. 2a). In the limit of large *C*_ind_ the membrane energy is dominated by the bending energy and we can approximate the membrane shape by a cylinder with radius 1*/*(2*C*_ind_) and length *L* = *A*_c_*C*_ind_*/π* (dotted line in Fig. 2a). The energy difference between the flat and the cylindrical shape is given by 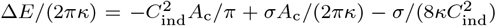. The cylindrical shape becomes energetically advantageous compared to the spherical cap if the induced spontaneous curvature exceeds a value of 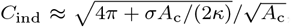, in the limit *σA*_c_*/κ* ≫ 1. This simple approximation shows that, as the protein density or equivalent of the induced spontaneous curvature increases, the height of the protrusion increases linearly with *C*_ind_. We notice that the membrane shape will transition from a spherical cap shape to a cylindrical shape at the threshold of 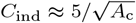 (green marker in Fig. 2a).

**Figure 2:**
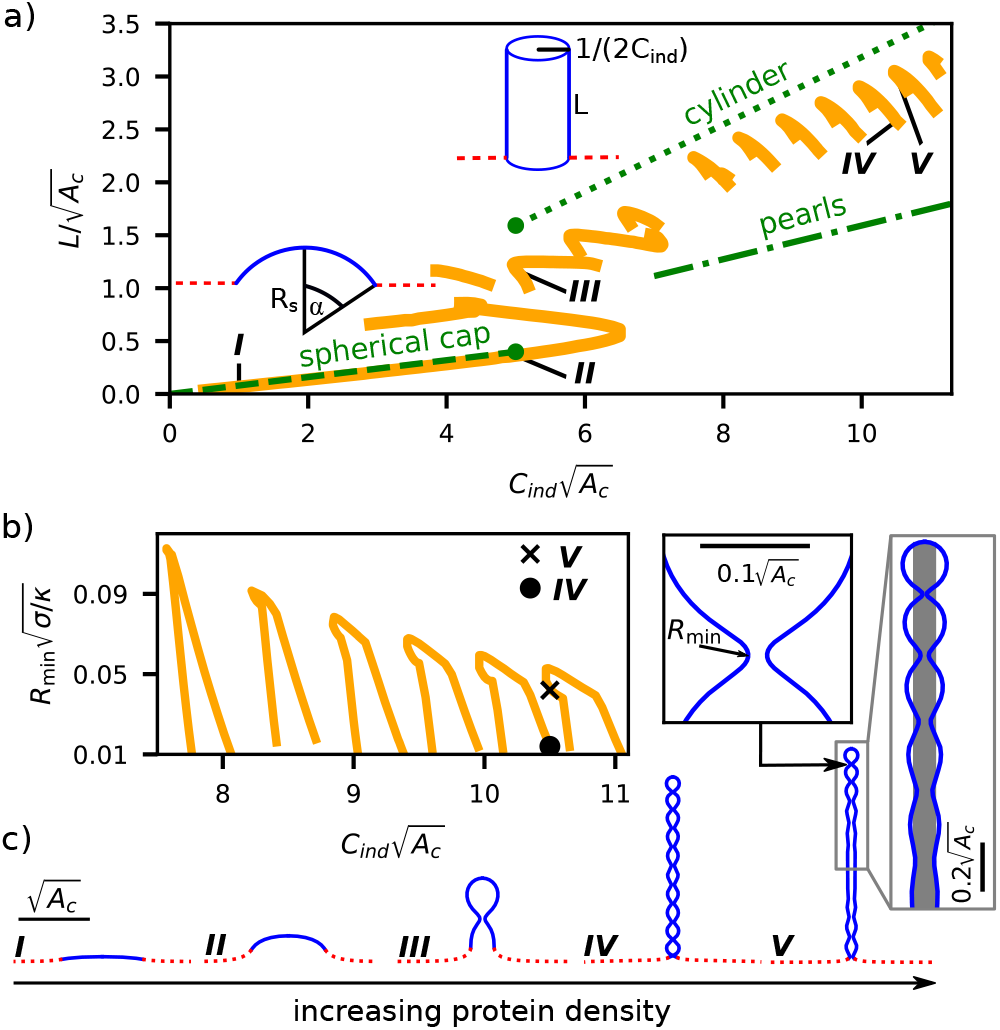
a) The scaled protrusion height *L* is shown as a function of the induced spontaneous curvature *C*_ind_. The results from the shape equations (Eqs. 7) are shown by orange solid lines. The analytic approximations for a spherical cap (dashed), a cylinder (dotted) and a string of beads (dash-dotted) are shown as green lines. At the transition point (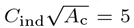, green marker) the energy of a spherical cap and a cylinder are equal. b) The minimal neck radius *R*_min_ as a function of the induced spontaneous curvature for tubular shapes with 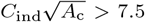. c) Five membrane shapes are shown, where the crowded region is illustrated by the solid blue line, while the protein free region is shown as a red dashed line. The scale bar indicates the length 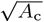 that is set by the area of the crowded domain. The inset to the right of shape ***V*** shows an undulating tubular shape together with a cylinder with radius 1*/*(2*C*_ind_) in grey. The inset to the left shows the smallest membrane neck along the tubular shape, with 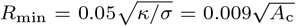, which corresponds to *R*_min_ = 40nm in dimensional units. Hence, *R*_min_ is still much larger than the membrane thickness.

We now turn to the energy minimizing shapes and scale all lengths by 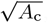, *i*.*e*., 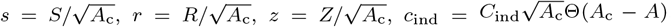 with the Heaviside function Θ, *ψ*(*S*) → *ψ*(*s*), *a* = *A/A*_c_, and 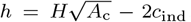. Applying the Euler Lagrange formalism, we determine the stationary shapes of the functional 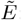:

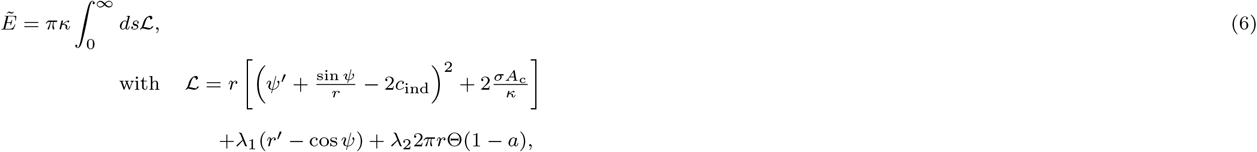

where the derivative with respect to *s* is indicated by a prime, *d*()*/ds* = ()^*′*^. In Eq. 6, the Lagrange multiplier function *λ*_1_ constrains the geometric relation between the azimuthal angle *ψ* and the radius *r*. The Lagrange multiplier *λ*_2_ enforces a constant area at the crowded domain. The energy minimizing membrane shapes are then described by the following set of differential equations (see SM):

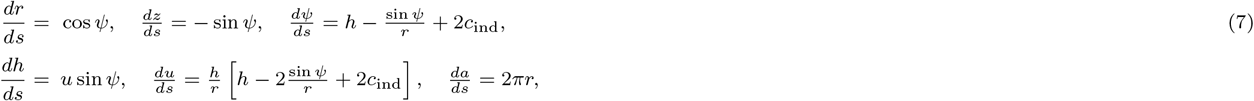

where *u* := *λ*_1_*/*(2*r*) is an auxiliary function that becomes equal to the scaled membrane tension at the outer boundary. The boundary conditions

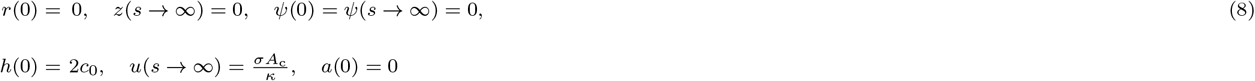

describe a membrane that is flat in the far field, while the mean curvature at the centerline (*s* = 0) is given by 2*c*_0_, where *c*_0_ is part of the solution of Eqs. 7. To solve Eqs. 7, we define the parameter *u*_0_ := *u*(*s* = 0) and treat the differential equation as an initial value problem where the parameters *c*_0_ and *u*_0_ are systematically varied such that the boundary conditions (Eq. 8) at the outer boundary are satisfied. Further details about the numerical implementation of Eqs. 7 can be found in the SM.

## Results and Discussion

In Fig. 2a the protrusion height *L* is shown as a function of *C*_ind_. *L* is defined as the height of the membrane in *Z*-direction above the center point *Z* = 0, *R* = 0 (Fig. 1 c). Similar to the analytic approximation presented above, we see a transition from flat shapes (shape ***I***) for small 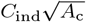 to elongated shapes (shape ***IV*** and ***V***) for large 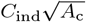. In the transition region around 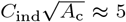, multiple stable shapes coexist, including shape ***II*** and ***III***. As we see in Fig. 2c, the shape of the tubulating membrane does not follow a cylindrical shape, instead the radius undulates around a value of 1*/*(2*C*_ind_) (grey area in shape ***V***), which is determined by the spontaneous curvature. The neck size of shape ***V*** is still much larger than the typical thickness of a lipid bilayer, highlighted by the inset. We note that even though the membrane has a shape similar to a string of beads, the protrusion height is significantly longer than if it would be comprised of spherical beads with radius 1*/C*_ind_, with *L* = *A*_c_*C*_ind_*/*(2*π*) (dash-dotted line in Fig. 2a). As the induced spontaneous curvature increases, Fig. 2a shows a series of branches. The tube length increases in a discontinous manner (Fig. 2a) between two consecutive branches. Each step originates from an additional bead along the tube (see shape ***IV*** and ***V***). Toward the ends of each branch the minimal neck radius *R*_min_ decreases as shown in Fig. 2b. As *R*_min_ decreases, the neck is constricted and eventually reach a size that is smaller than the thickness of the membrane. In this case the description of the membrane as a thin elastic sheet is no longer valid. We therefore limit the results shown in Fig. 2 to shapes with a neck size that is at least 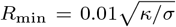, or equivalently *R*_min_ = 10nm. Hence, in all shapes considered here, the membrane neck is sufficiently wide to prevent membrane fission.

The membrane tubes we find here differ significantly from tubes caused by a point force, where the tube radius is determined by the ratio of the membrane tension and the bending rigidity showing only slight deviations from a cylindrical shape [14]. In contrast, tubes induced by crowding or by a change in the area-to-volume ratio by osmotic deflation [16] have qualitatively similar shapes, where the tube radius is determined by either the induced spontaneous curvature or the intrinsic membrane curvature. Despite the similar membrane shapes, there are large differences between these tubes. In the case of osmotic deflation, the area of a tube is not fixed and could in principle consume the entire surface of the original vesicle. In contrast, the total area of a tube induced by protein crowding is limited by the size of the crowded domain, and the spontaneous curvature is not fixed but is determined by the protein density.

As we follow the minimal energy shapes for increasing spontaneous curvature or equivalently for increasing protein density in Fig. 2a and Fig. 2c, we find shapes that are qualitatively similar to those recently found experimentally by Shurer et al. [20] on cells with glycocalyx polymers (Fig. 1b), including Ω-shapes/blebs (shape ***II***), tubes (shape ***V***) and pearls/beads (shape ***IV***). While several of these shapes have been theoretically described before in specific analytical limits [20], we here obtain the entire variety of shapes by the a single underlying physical principle, *i*.*e*. the interplay of the induced spontaneous curvature and the membrane tension, where all shapes are described by the same set of shape equations (Eqs. 7).

To relate the induced spontaneous curvature to the protein density *ρ* based on Eq. 5 we need to know the radius of the proteins *r*_p_and the bending rigidity *κ*. We set *r*_p_ = 2.1nm and *κ* = 10k_B_T in accordance with the values given by Stachowiak et al. in an experimental study on the tubulation propensity of GUVs covered with density packed intrinsically unstructured proteins. In addition, we assume a crowded area of *A*_c_ = 50*µ*m^2^. In Fig. 3 the protrusion height *L* is shown as a function of the density *ρ*. To define the onset of tubulation, we require that the protrusion height reaches 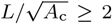, which is satisfied for 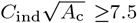 (Fig. 2a). We obtain a threshold density for tubulation of (*ρ/ρ*_max_)_tube_ ≈ 0.19. A smaller crowded domain require larger protein densities to generate tubulation and *vice versa, e*.*g*. (*ρ/ρ*_max_)_tube_ = 0.12 for *A*_c_ = 500*µ*m^2^ or (*ρ/ρ*_max_)_tube_ = 0.28 for *A*_c_ = 5*µ*m^2^. Stachowiak et al. investigated experimentally the relation between the propensity to tubulate and the protein density [26]. They found the fraction *f* of GUVs that form tubes to increase with increasing protein coverage (experimental data reproduced in the inset in Fig. 3). In these experiments the GUVs and consequently also the crowded domain vary in size, where only a fraction of the GUVs exhibits a crowded area with a protein density that together are large enough to induce tubulation. From the experimental measurements it is known that the crowded domain comprises about 15% of the total GUV surface [26], in addition, we assume that the GUV radii are equally distributed up to a maximal radius *R*_max_. By fitting the experimental data to the theoretically expected fraction of vesicles that exhibit tubulation (see SM) we obtain *R*_max_= (10 ± 3)*µ*m, which agrees well with a typical GUV size. The comparison between our theoretical model and the experimental data indicates that to observe tubulation both the protein density and the crowded area must in combination exceed a critical value.

**Figure 3:**
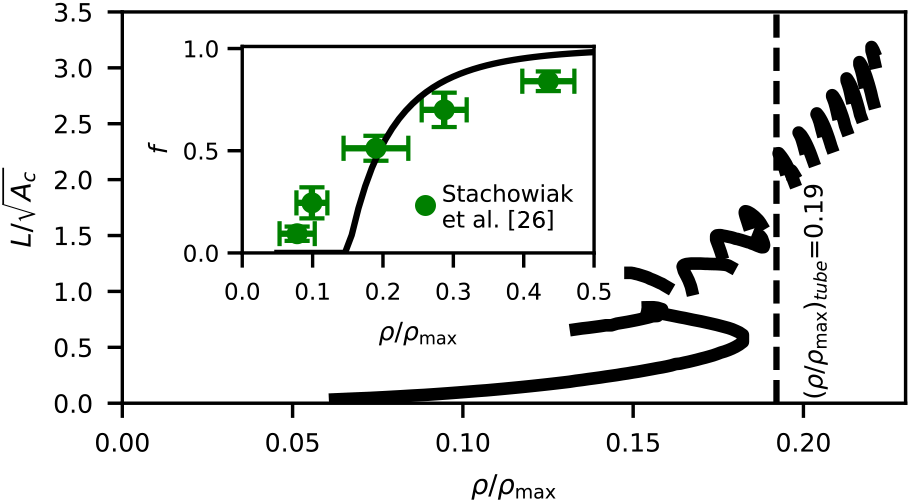
The protrusion height *L* is shown as a function of the protein density *ρ*, where *C*_ind_ (Fig. 2a) and *ρ/ρ*_max_ are related via Eq. 5. The inset shows the fraction *f* of GUVs that form tubes in the experimental study by Stachowiak et al. [26] (circular markers) and according to our theoretical model (solid line).

So far we have considered a fixed value for the scaled membrane tension. We now discuss the dependence of the membrane shape on the membrane tension *σ* for a fixed *C*_ind_. In Fig. 4 we show *L* as a function of the scaled membrane tension *σA*_c_*/κ* for three different values of 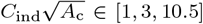. The inset in each subfigure shows the membrane neck radius *R*_min_. If the membrane shape does not exhibit an indentation (*e*.*g*., shape ***I***), *R*_min_ is defined as the radius at the outer edge of the crowded domain. As in Fig. 2, we only show membrane shapes with a neck radius of 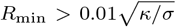, *i*.*e*. the neck is larger than the membrane thickness and the thin sheet description of the membrane is valid. The three shapes that are indicated by roman numbers are also shown in Fig. 2. For the lowest value, 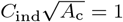, a monotonic decrease of *L* with increasing membrane tension is observed, *i*.*e*., the membrane flattens (shape ***I***).

**Figure 4:**
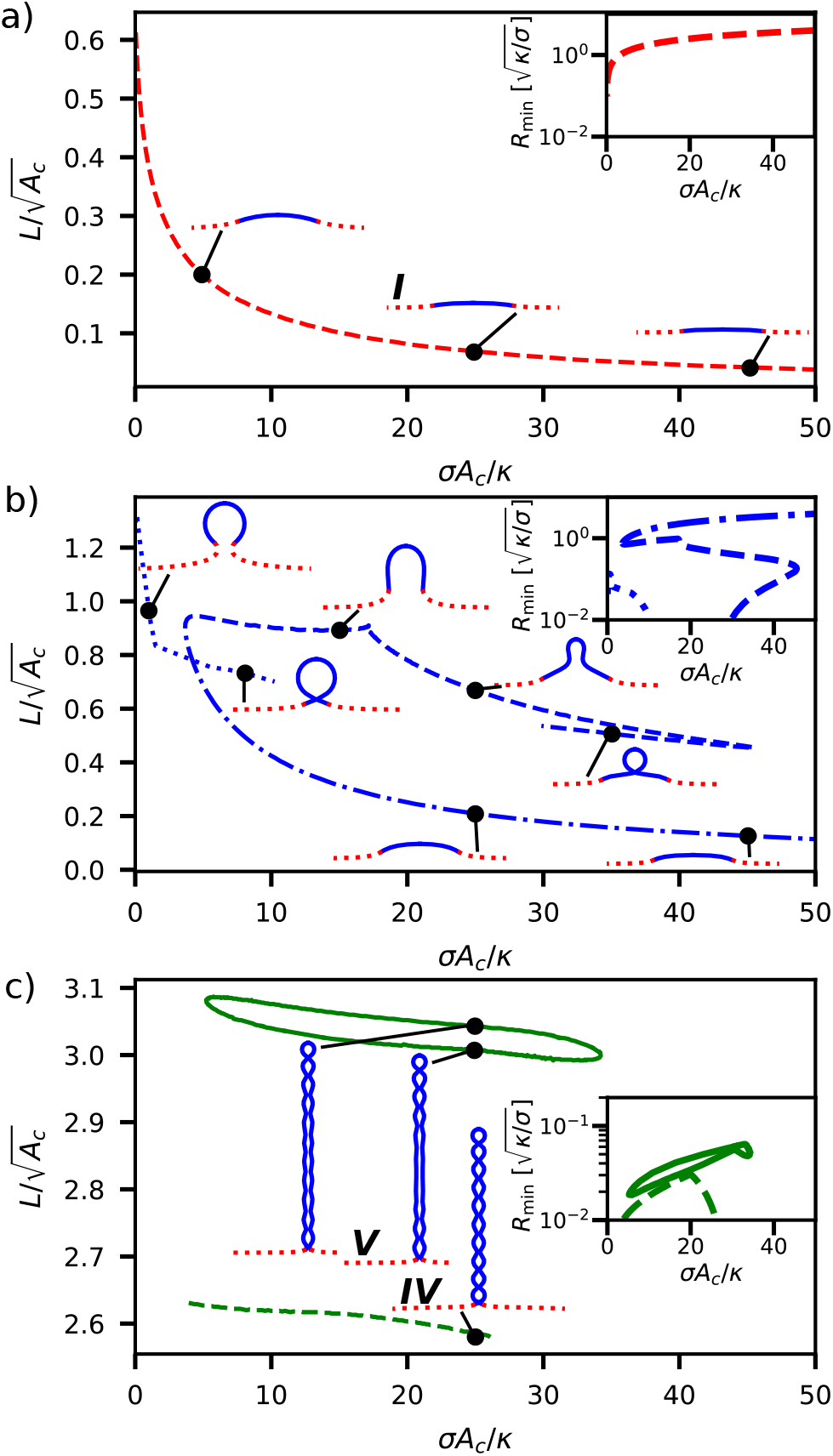
The scaled protrusion height *L* is shown as a function of the scaled membrane tension *σA*_c_*/κ* for different induced spontaneous curvatures *C*_ind_. The inset in each subfigure shows the membrane neck radius *R*_min_. The three shapes that are indicated by roman numbers are the same shapes shown in Fig. 2.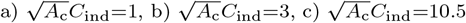.

Increasing the spontaneous curvature to 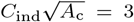, we find a more complex behavior with three different classes of membrane equilibrium shapes (shown in Fig. 4b as dashed, dashed-dotted, and dotted lines, respectively) that can coexist. The lower line (dashed-dotted line) corresponds to flattened shapes that minimizes the tension energy. The shapes in the upper dashed line show a narrower protrusion. For *σA*_c_*/κ <* 12 there is a third class of equilibrium shapes that is associated with spherical membrane shapes. These shapes do not lead to tubulation and the neck becomes narrower as *σA*_c_*/κ* increases. We speculate that a further increase of the membrane tension (*σA*_c_*/κ >* 12) would lead to an overlap of opposing sides of the neck and subsequent membrane fission. Membrane scission by protein crowding, was observed experimentally by Snead *et al*. [34], where the presence of unstructured proteins (an epsin1 N-terminal homology domain) on the membrane was sufficient to induce vesicle formation even in the absence of specialised proteins that facilitate membrane scission. For an even larger spontaneous curvature, 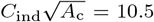, elongated tubes are formed. While we can observe a closed line for shapes with eleven connected beads along the tube (shape ***V***), shapes with only ten beads (shape ***IV***) are limited by a constriction of the membrane neck for both low and high membrane tensions.

The minimal theoretical model presented here shows that steric repulsion between membrane-associated proteins and membrane tension can explain a variety of membrane shapes observed in experiments, providing a quantitative framework for understanding membrane remodeling induced by protein crowding. In Fig. 2 and Fig. 3 we have shown that multiple equilibrium membrane shapes can coexist: flat, spherical and tubular shapes. Our results highlight protein crowding as a versatile mechanism for membrane shape regulation, which is a process vital to cell functionally by compartmentalising or connecting cellular organelles.

## Acknowledgement

We thank Dr. Padmini Rangamani for stimulating discussions and input to this manuscript. The work has been enabled by funding from the Research Council of Norway (Project Grant 263056).

